# The contribution of global mountains to the latitudinal diversity gradient

**DOI:** 10.1101/2020.07.04.188227

**Authors:** Elkin A. Tenorio, Paola Montoya, Natalia Norden, Susana Rodríguez-Buriticá, Beatriz Salgado-Negret, Mailyn A. Gonzalez

## Abstract

The latitudinal diversity gradient (LDG) is widely attributed to be the result of factors such as time, area, and energy. Although these factors explain most of the variation in lowlands, they fail in mountainous systems, which are biodiversity hotspots that may contribute meaningfully to the strength of the pattern following different evolutionary pathways. However, because lowlands cover the largest portion of the total land, they may have overshadowed the contribution of mountains to the LDG, but no study has addressed this issue in previous macroecological analyses. Here, we propose that the LDG shows a stronger trend in mountain ranges due to their high species turnover, in spite of covering less than one third of the Earth’s land. Using the geographical information for ∼22000 species of terrestrial vertebrates, we show that worldwide mountains harbor the 40% of the global diversity, and when taking into account the area effect, we quantified that mountains harbor close to double the species inhabiting lowlands per unit area. Moreover, when we evaluated the LDG after accounting for area size, we found that species richness increased faster towards the Equator and was better predicted by latitude in mountains than in lowlands. Our findings challenge previously well-supported hypotheses that predict that those regions with greater area, time and energy accumulate more species richness, since mountains are geologically younger, exhibit less energy, and cover smaller areas than lowlands. Hence, mountains represent a paradox, which invites to reevaluate hypotheses regarding macroecological and evolutionary processes driving species diversity gradients.

## Introduction

The increase in species diversity towards the Equator is one of the most consistent and well-known patterns in ecology (1, 2), yet its underlying drivers remain elusive. Area, energy, and time are among the factors that best explain the latitudinal diversity gradient (LDG), as more species are accumulated in older and larger areas with higher productivity, where diversification is promoted by low extinction rates or frequent speciation events (1–4). Mountains, however, represent a paradox, as they may harbor exceptional levels of biodiversity in small areas (5, 6), are characterized by low levels of productivity in high elevations (7), are often geologically younger than surrounding lowlands, and have had less time to be colonized by clades that could have undergone subsequent diversification (8). In fact, several mountain ranges such as the Andes, the Eastern Arc mountains, and the Indo-Pacific mountainous islands are well recognized as biodiversity hotspots (9–11).

Previous attempts to relate regional species richness with contemporary climate, net primary productivity, and topography have failed to explain the high levels of diversity observed in mountains (5, 12). This suggests that the relative influence of evolutionary processes determining spatial patterns of species richness are different regarding lowlands. The contrasting patterns of species turnover (or beta-diversity) between landforms could be a consequence of such differential influence. Though species segregation is higher in tropical latitudes, it is strikingly prominent in tropical mountains, likely as a result of the dramatic change in abiotic conditions between adjacent elevational thermal belts, which contrasts with the less extreme zonation in temperate regions (5). This high zonation in tropical mountains facilitates the strong replacement of almost entire communities over short geographical distances. Considering the smaller area of mountains, the species replacement along slopes of mountain ranges may result in a greater capacity to harbor more species per unit area and, in turn, generate a greater species packing when compared to the lowlands. Thus, a higher species turnover in tropical mountains might have an additive effect on the inherent increase of diversity towards low latitudes, leading to a more pronounced LDG. Although the association between latitude and beta-diversity in mountains has been previously noticed (13, 14), the magnitude to which it may generate an additive effect causing a difference in the LDG in mountains has not yet been explored.

Here, we hypothesize that latitudinal gradients are much stronger in mountains than in lowlands as a result of species packing and turnover. Addressing this issue has been challenging because lowland areas are geographically larger, and when the effect of area is not accounted for, the importance of mountains in determining the overall patterns of species diversity may be overshadowed. We quantify the contribution of mountains to worldwide species richness in three groups of terrestrial vertebrates to test whether the LDG is steeper in mountains than in lowlands after accounting for area size.

## Results and Discussion

To address this issue, we first evaluated the total contribution of mountains to global patterns of bird, amphibian, and mammal species richness (ca. 22,000 species). Using public data on the global distribution of each of these groups, we calculated the proportion of each species’ distribution in mountains vs. lowlands based on two alternative definitions of mountains: 1) areas above 700 m, a consensus on elevational limits used in several studies, and 2) areas with “high ruggedness,” a criterion based upon elevational differences within a given area (6). A species was considered mountainous if the proportion of its distribution in mountains was higher than a given threshold (*T*), such that a threshold of *T*=0.5 meant that a species was considered mountainous when at least 50% of its distribution range was in mountains. We quantified global species richness in each group of vertebrates using a *T*=0.5 and examined the robustness of our results by calculating a confidence interval based on *T*=0.3 and *T*=0.7 (see details in Methods).

Worldwide, we found that mountains were disproportionately species-rich compared to lowlands when accounting for area. Based on the elevation-driven definition of a mountain and a threshold of *T*=0.5, mountains occupied only 28.5% of emerged lands, yet they harbored as much as 39% of the world’s diversity of terrestrial vertebrates. This net estimation of number of species translates to 1.6 times more species per unit area than lowlands (Fig. 1B, Table 1). These results held even when using *T*=0.7, a highly conservative threshold for considering a species as mountainous, where mountains harbored 31.5% worldwide diversity (Table 1). By evaluating groups separately, mountains held 1.3, 1.4 and 2.4 more birds, mammals and amphibians species respectively per unit area than lowlands (Figs. 1C,D,E, Table 1). Patterns were stronger when defining mountains based upon ruggedness, where mountains represented only 18.5% of emerged lands, but still containing the 39% of terrestrial vertebrates biodiversity. In other words, mountains held 2.9 times more richness per unit area than lowlands (*T*=0.5, SI Appendix, Fig. S1 and Table S1). Our findings contrast with those of Rahbek et al (5), who recently reported that mountains shelter 87% of terrestrial vertebrate species. We believe this value corresponds to an overestimation, as they included in their calculations any species occupying tangential areas in mountains. In this sense, our estimates are more conservative, as we considered a species to be mountainous only when a significant proportion of its distribution felt in mountainous landscapes.

**Table 1.**
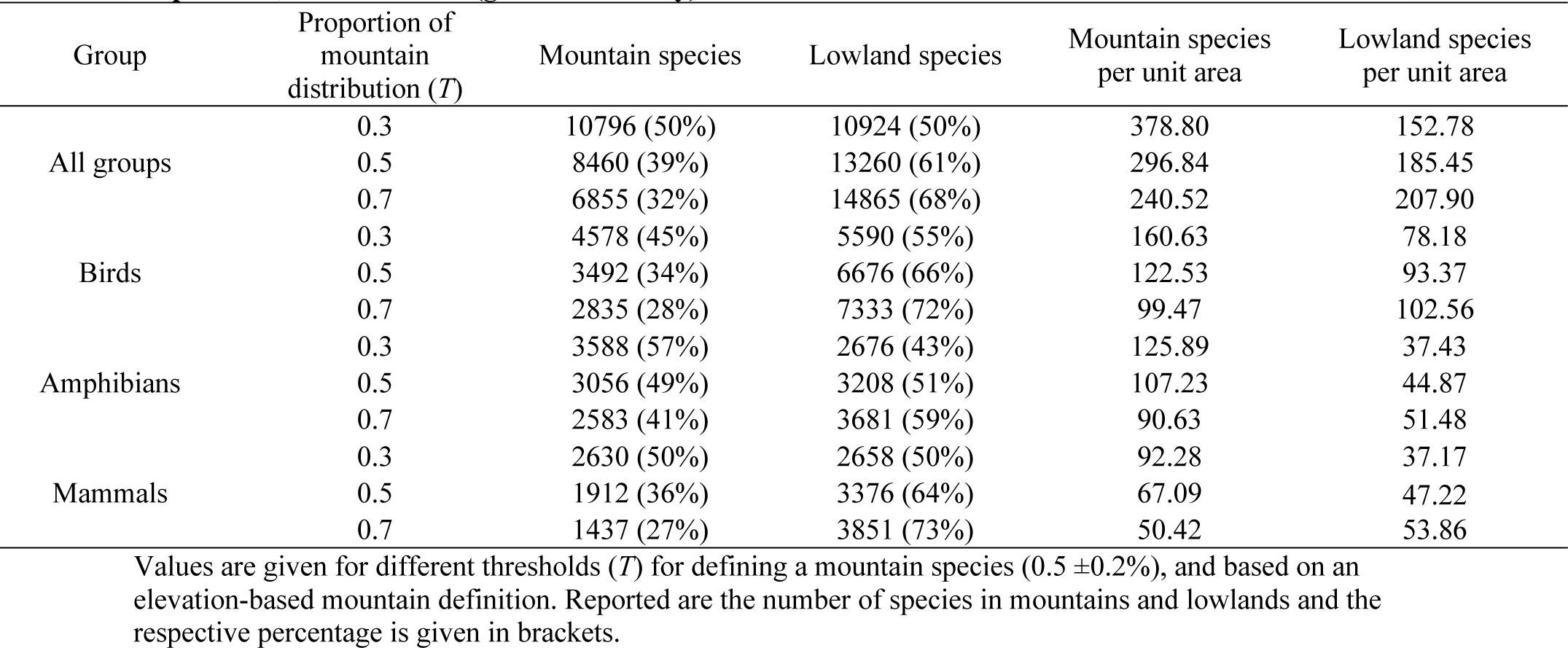
Estimation of mountain and lowland contribution to the total species richness of birds, amphibians, and mammals (gamma diversity).

| Group | Proportion of mountain distribution (*T*) | Mountain species | Lowland species | Mountain species per unit area | Lowland species per unit area |
| --- | --- | --- | --- | --- | --- |
| All groups | 0.3 | 10796 (50%) | 10924 (50%) | 378.80 | 152.78 |
|  | 0.5 | 8460 (39%) | 13260 (61%) | 296.84 | 185.45 |
|  | 0.7 | 6855 (32%) | 14865 (68%) | 240.52 | 207.90 |
| Birds | 0.3 | 4578 (45%) | 5590 (55%) | 160.63 | 78.18 |
|  | 0.5 | 3492 (34%) | 6676 (66%) | 122.53 | 93.37 |
|  | 0.7 | 2835 (28%) | 7333 (72%) | 99.47 | 102.56 |
| Amphibians | 0.3 | 3588 (57%) | 2676 (43%) | 125.89 | 37.43 |
|  | 0.5 | 3056 (49%) | 3208 (51%) | 107.23 | 44.87 |
|  | 0.7 | 2583 (41%) | 3681 (59%) | 90.63 | 51.48 |
| Mammals | 0.3 | 2630 (50%) | 2658 (50%) | 92.28 | 37.17 |
|  | 0.5 | 1912 (36%) | 3376 (64%) | 67.09 | 47.22 |
|  | 0.7 | 1437 (27%) | 3851 (73%) | 50.42 | 53.86 |

Values are given for different thresholds (*T*) for defining a mountain species ( $0.5 \pm 0.2\%$ ), and based on an elevation-based mountain definition. Reported are the number of species in mountains and lowlands and the respective percentage is given in brackets.

**Fig. 1.**
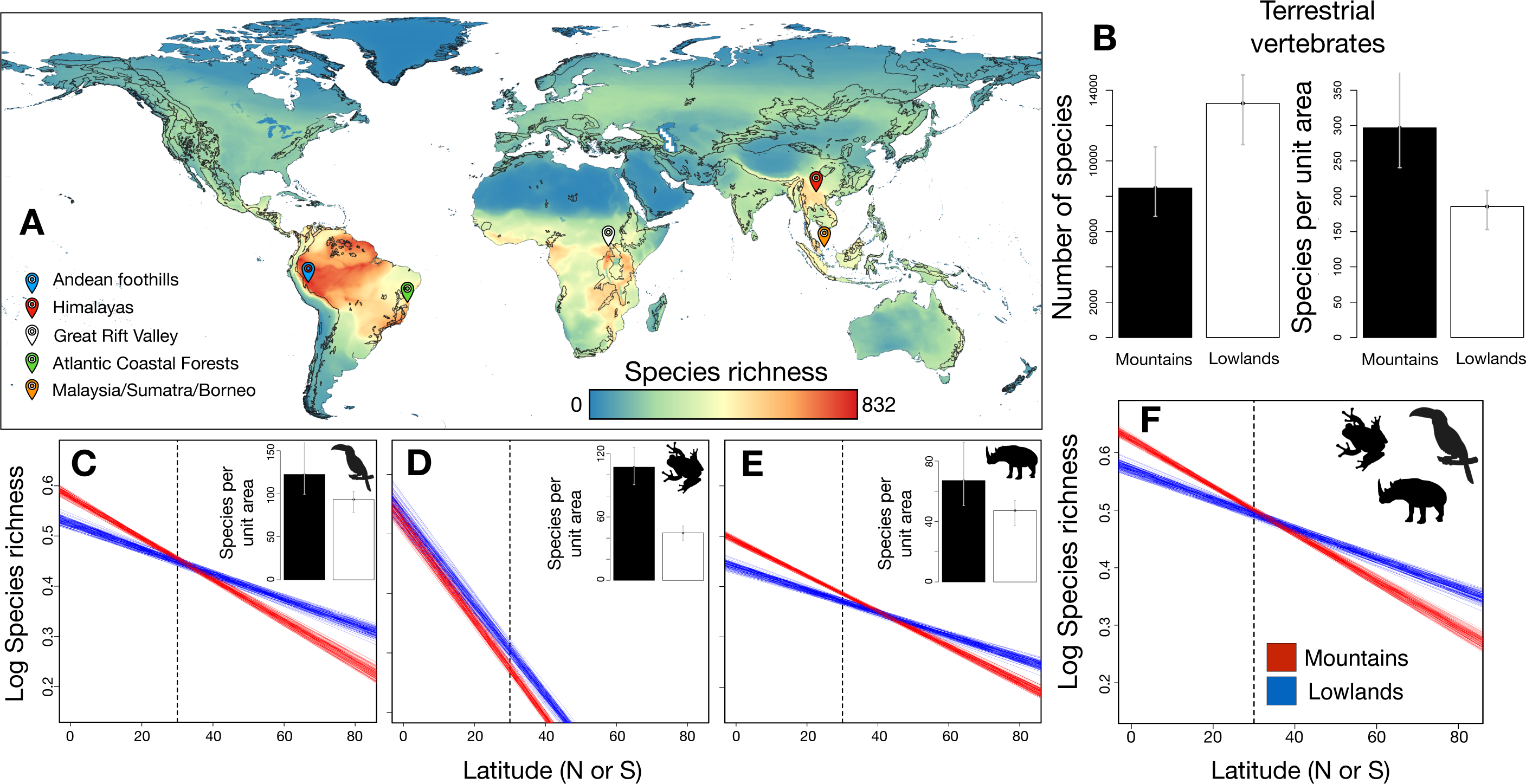
Global patterns of species richness in mountains vs. lowlands. (A) Species richness map of the three groups of terrestrial vertebrates (birds, mammals, and amphibians). Mountains regions are outlined by a black line following the inventory of the World’s mountains from the Global Mountain Biodiversity Assessment (GMBA, http://www.mountainbiodiversity.org). Excepting the Amazon, highest values of species richness are associated mainly with tropical mountain systems, as exemplified by the color marks highlighting five mountain ranges (Andean foothills, Himalayas, Great Rift Valley, Atlantic Coastal Forest and mountains in Malaysia, Sumatra, and Borneo). (B) Differences in the contribution of mountains (black bars) and lowlands (white bars) to the total species richness combining the three groups of terrestrial vertebrates based on an elevation criterion for defining mountains. The bars in (B) represent the total number of species and the number of species per unit area for all three groups, and separately for (C) birds (D) amphibians, and (E) mammals. In each panel, bar height corresponds to estimations using a *T*=0.5 and intervals (gray vertical line) correspond to values generated by varying *T*, the threshold for defining a species distribution as being in mountains vs. lowlands (*T*= 0.5±0.2%, *T*=0.3% and *T*= 0.7% – values in Table 1). (F) Patterns of species richness along the latitudinal gradient for mountains (red) and lowlands (blue) when area is controlled for and using an elevation criterion to define mountains; analyses were performed for all vertebrates (F), and separately for (C) birds (D) amphibians, and (E) mammals. Trend lines were extracted from 100 linear models constructed using 1000 random pixels for both mountains and lowlands. The dotted black line represents 30° latitude, the subtropical-temperate limit where the average richness per latitude point is higher in mountains. Silhouettes were taken from PhyloPics (credits to Will Booker, FJDegrange, and Zimices).

Heightened diversity in mountains could be explained by topographic and climatic heterogeneity (13), which act as barriers to dispersal favoring the segregation of species with restricted dispersal abilities, thereby magnifying beta-diversity, which in turns enhances gamma diversity (13, 15). This species segregation also reflects the effects of historical and evolutionary forces that have promoted lineage accumulation in mountain ranges (16, 17). Indeed, there is evidence that mountains have greater diversification rates than lowlands (18), likely resulting from differences in the relative influences of processes occurring in each of these landforms. For example, tectonic uplift probably offered new and unexplored arenas, ready to be colonized by lowland clades or temperate migrants (19), which in turn may undergo speciation through vicariance, facilitated by niche conservatism or ecological opportunity in new high elevation environments (20–22). High diversification rates in mountains could also be driven by lower extinction rates, as species may be less vulnerable to environmental changes given that they need to disperse shorter distances up or down slope to escape unsuitable areas compared to lowlands (17). Overall, the relative contribution of speciation versus extinction rates to differences in diversification rates between mountains and lowlands needs to be evaluated in future studies.

Given the extraordinary species richness of mountains, we tested whether the strength of the latitudinal gradient in mountains is different from that in lowlands, when we take into account the area. We evaluated the relationship between species richness and latitude in each landform for all three vertebrate groups combined and separately. Species richness was estimated at a finer pixel resolution (10km x 10km; Fig. 1A) than previous macroecological works (*e*.*g*. ∼110km x 110km). This enabled us to minimize the mixing of mountain and lowland areas within the same pixel, and to reduce mountain underrepresentation resulting from their low extension (see Methods). To control for spatial autocorrelation, we implemented a Monte Carlo procedure, where, for each of 1000 iterations, we randomly selected 1000 pixels located in mountains and 1000 pixels located in lowlands across the globe, and estimated the slope (β) and coefficient of determination (*R*^*2*^) of the linear regression predicting species richness from latitude.

Using the elevation-based definition of mountains, we found a negative relationship between species richness and latitude for both landforms, but slopes (β) and coefficients of determination (*R*^*2*^) were significantly higher for mountains (ANOVA: *P* < 0.001; Fig. 1F, SI Appendix, Fig. S2 and Table S2). In other words, species richness increased faster towards the Equator and was better predicted by latitude in mountains than in lowlands. Remarkably, the latitudinal point where the slopes of the two models crossed, i.e., where species richness becomes higher in mountains than in lowlands towards the Equator, roughly corresponded to the subtropical-temperate latitudinal limit (especially in birds and mammals). A stronger latitudinal gradient in mountains supports the untested hypothesis put forth by Simpson (14), who claimed that “where there are latitudinal gradients, these are additive with topographic gradients, the two accounting for most of the pattern.” Our findings also concur with patterns reported by Fjeldså (23), where mountains were particularly species-poor in temperate latitudes, especially above 20-40° latitude. When using the ruggedness-based definition of mountains, the slopes describing the LDG were apparently parallel and never crossed, although they were statistically steeper for lowlands than for mountains (*P* < 0.001; SI Appendix, Fig. S3 and S4). However, as in the elevation-based analysis, latitude explained a higher fraction of the variance in species richness in mountains than in the lowlands (*P* < 0.001; SI Appendix, Table S3).

The fact that differences between mountains and lowlands were more conspicuous using the elevation-based definition of mountains suggests that elevation is more strongly related to the underlying processes determining the LDG in mountains than topographic complexity alone. Specifically, variables correlated with elevation, such as oxygen concentration and temperature, are likely to limit vertebrate physiological responses along elevation gradients but not across topographical heterogeneity. Indeed, the fact that mountains showed higher diversity than lowlands below the subtropical-temperate latitudinal limit suggests that there might be region-specific processes shaping patterns of vertebrate species richness in the tropics. This finding fits with the idea that “mountain passes are higher in the tropics” (24, 25), which provides a powerful mechanistic explanation for why conditions in tropical mountains generate an additive effect that magnifies the latitudinal gradient of species richness. D. H. Janzen (24) suggested that topographical barriers are more efficient in tropical than in temperate latitudes because tropical species are expected to have narrower thermal tolerances in response to higher thermal stability in tropical mountains. Narrow thermal tolerance increases the cost of dispersal across thermal belts, which may reduce gene flow between tropical populations and eventually may facilitate allopatric speciation (21, 25, 26).

When evaluating each group separately, mammals and birds exhibited similar patterns to the overall trend (*P* < 0.001; Figs. 1C and E) but in amphibians the rate of increment in species richness towards the Equator was similar between landforms (*P* < 0.001; Fig. 1D, SI Appendix, Fig. S2 and Table S2). This finding could result from differences in dispersal abilities and contrasting physiological features between ectotherms (amphibians) and endotherms (birds and mammals), which in turn is expected to affect their spatial distribution. Unlike endotherms, the major restriction for amphibian distribution at high elevations is not determined by thermal tolerance but rather by humidity and ultraviolet radiation (27, 28). If so, changes in temperature associated with mountain topographic heterogeneity should be more efficient barriers for mammals and birds than for amphibians, therefore explaining the different patterns in the LDG observed between ectotherms and endotherms.

As mountainous regions harbor most of the range-restricted species occurring worldwide (11, 17), one may wonder whether our results were affected by the predominance of this group of species. To address this issue, we evaluated the relationship between species richness and latitude by partitioning species into range size quartiles (12, 29). Patterns of species richness along the latitudinal gradient for range-restricted (narrowest ranging 25%) and widespread (widest ranging 25%) species were alike (SI Appendix, Fig. S5), indicating that our findings were not biased by differences in species’ spatial distribution between landforms. We also evaluated whether our results were driven by region-specific patterns which could be generating the global pattern. For instance, the Andes Cordillera is well-known to concentrate high levels of diversity, especially of range-restricted species, compared to other large mountain systems like the Himalayas or any other system in the Eastern Hemisphere (11, 30). We therefore analyzed our data by accounting only for species occurring either in the New World or in the Old World, yet both regions showed congruent results (Fig. 2, SI Appendix Fig. S6), supporting the idea that our findings correspond to a generalized trend across the globe.

**Fig. 2.**
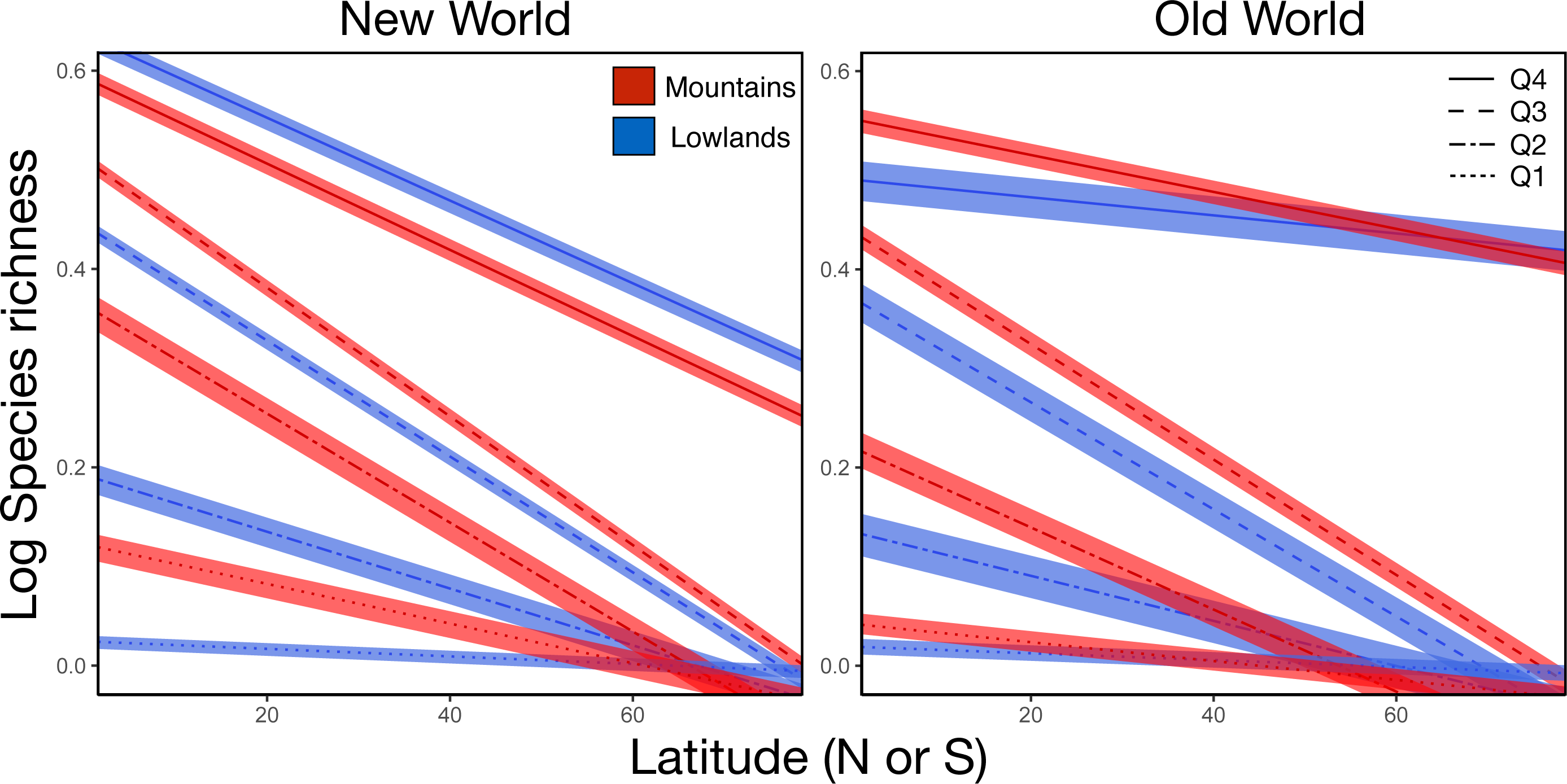
Species richness patterns by range size quartiles of terrestrial vertebrates richness (birds, amphibians, and mammals) in the New World and the Old World. Analysis was performed using a resample method to control by area and using an elevation criterion to limit mountains. Species were divided in four geographic range size quartiles, from the narrowest ranging species (1^st^, 0-25%) to the widest ranging (4^th^, 75-100%), and the line pattern represents each quantile: solid line (1^st^ quantile), dashed line (2^nd^ quantile), dash-dotted line (3^th^ quantile), and dotted line (4^th^ quantile). Envelopes around each line show the 95% confidence interval generated by extracting the trend lines from 1000 linear models based on a random sample of 1000 pixels for mountains and lowlands separately for mountains (red) and lowlands (blue) in each quantile and continent.

Overall, we show that mountains harbor a higher diversity per unit area and exhibit a steeper LDG than lowlands. Our results have critical insights for the understanding of the LDG, one of the most striking patterns in the distribution of biodiversity. In particular, the fact that species richness of terrestrial vertebrates per unit area in mountains surpassed that of lowlands challenges the time-area hypothesis, where older and larger areas with historically higher energy availability are more likely to accumulate greater diversity (1, 31, 32). Although we do not provide an explicit test of this hypothesis, our results contrast with its predictions in different ways. First, mountains contributed nearly 40% of the world’s gamma diversity, despite covering less than one third of the available terrestrial area. Second, mountains tend to have lower energy in terms of primary productivity than lowlands. Third, available time for colonization and diversification in mountains is likely to be lower than in surrounding lowlands, not only due to relatively recent orogenic processes, but also because glacial cycles have recurrently erased biotas of high elevations in tropical and temperate mountains (33). It is therefore essential to integrate the effect of time since colonization in future studies evaluating LDG, because it may influence the net cumulative diversity in mountains and lowlands. Doing so would require geological and paleoclimatic information combined with data on orogeny, which are not available yet. Until then, our results shed reasonable doubt on the explanatory power of the time-area hypothesis.

If the relative influence of diversification processes associated with spatial heterogeneity, dispersal mechanisms, and reproductive isolation varies from lowlands to mountains (17, 30, 33), evaluating these two landforms without any distinction may obscure the underlying processes determining global patterns of species richness. We therefore need to recognize a possible effect of region and control for the effect of area to avoid overshadowing the influence of mountains in macroecological analyses. When considering that tropical mountains are among the most diverse yet threatened systems by human activities and climate change worldwide (34, 35), our results have important implications in conservation biology. Given the exceptional levels of species richness in mountains, if we want to understand the processes that generate *worldwide* diversity, we must first understand the processes that generate overall *mountain* diversity.

## Methods

### Definitions of mountains

We selected two criteria proposed in the literature to define mountains: one based on an elevational limit threshold and another based on ruggedness (6). Based on a landmass grid of 10 km x 10 km, we classified each pixel as either mountain or lowland following each of these two criteria. The elevation criterion defined mountains as areas with elevation above 700 masl. Using this elevation limit and creating a mask based on a digital elevation model (SRTM Digital Elevation Database v4.1: http://www2.jpl.nasa.gov/srtm/), we found that the global area of mountains represented 28.5 % of the total land area. For ruggedness, we used the raster layer of mountains generated by Körner et al. (6). Ruggedness is defined as the maximal difference in elevation among neighboring grid points, where mountains were determined as pixels with at least 200 meters of ruggedness. This definition, however, may include isolated pixels that are not part of mountain ranges, usually at low elevations. To correct for this potential bias, we considered only pixels with an elevation above 200 meters (around 30% of pixels were discarded). Under this criterion, the global area of mountains represented 18.5% of total land area.

Both definitions show different landscape attributes that impact particular ecological and physiological traits (36). A classification of mountains based on elevation alone inflates the influence of variables like changes in air pressure, oxygen concentration, temperature, and UV radiation along elevation gradients, all of which demand physiological mechanisms that constrain animal distribution at high altitude. The downside is that thermal belts are not comparable for all latitudes and mountain systems, because features such as air currents, distance to the ocean, and topography also influence abiotic variables. Ruggedness, on the other hand, captures topographical complexity, but includes rough terrain at low elevations and excludes flat land at high elevations (e.g., the Andean plateaus in Colombia and Bolivia, the Rocky Mountains in the USA, or the Tibetan Plateau). Thus, ruggedness does not correlate with physical factors interacting with vertebrate physiological traits. Furthermore, ruggedness acts as a proxy of slope, which is more important in young mountain systems than in older ones, where erosion may have worn away steep slopes. For these reasons, we expected each definition to show different results given their features. For instance, defining mountains based on elevation may reflect climatic restrictions on species distribution, whereas doing so based on ruggedness emphasizes how physical attributes interact with species dispersal across barriers in mountainous areas. When evaluating the map congruence between both definitions, we found a match in 77% of the pixels, where the remaining 23% corresponded mainly to areas below 3000 masl. A map illustrating discrepancies between mountain areas according to both definitions in shown in SI Appendix, Fig. S7.

Although defining mountains as areas above 700 masl may be seen as arbitrary, any other threshold chosen would be as arbitrary as this one. This issue has been discussed in detail (37), and although different elevation limits have been used to differentiate mountains from lowlands, a limit of 1000 masl is the one most frequently used. Nonetheless, it has also been recognized that several mountain systems occur at elevations below 1000 masl in tropical areas (23), between 500 and 700 masl (37). We therefore present our results using the elevation thresholds of 700 masl and quantified the deviation of our estimates based on elevation limits of 500 and 1000 masl (see next section).

### Quantifying global mountain diversity

To evaluate the contribution of mountains to worldwide species richness in terrestrial vertebrates, we assembled a comprehensive database which maps species distribution for birds, amphibians and mammals at the finest resolution ever performed at the worldwide scale (10 km x 10 km). To do so, we quantified the extent to which the distribution range of each species overlaps with the masks for mountain areas obtained using the two definitions of mountains, respectively. We used expert distribution maps for birds (BirdLife International http://www.birdlife.org/datazone/home, accessed October 2014), amphibians, and mammals (IUCN Digital distribution maps http://www.iucnredlist.org/technical-documents/spatial-data, accessed October 2014). For birds, we only quantified the distribution of breeding areas of migratory species and excluded 238 marine species belonging to 10 families (Alcidae, Procellariidae, Spheniscidae, Stercorariidae, Laridae, Diomedeidae, Oceanitidae, Hydrobatidae, Sulidae, and Phaethontidae).

An issue pointed out in several macroecological studies at regional and global scales is the fact that expert maps for some species may have coarse limits and include zones outside the known distribution (for example, valleys or mountain peaks) (38). To reduce this bias and partially solve this problem at the scale and resolution of our study, especially in the case of birds, we constrained distribution maps to the known elevation range of each species, setting minimum and maximum elevation according to information in the Handbook of Birds of the World (39). We rasterized each species map to a 10 km x 10 km resolution using a WGS84 coordinate system. In some cases, mammals and amphibian species showed distribution ranges that were not detectable at the resolution used. In these specific cases, we rasterized their maps to 1 km x 1 km. In total, we used 21,741 species (10,186 birds, 6,266 amphibians, and 5,289 mammals). All procedures were undertaken using the libraries maptools (40) and raster (41) in R Software (42).

For each species and each mountain definition, we quantified the proportion of their distribution that overlapped with mountains using the rasterized maps separately (ranged from 0 for species showing a distribution entirely in the lowlands to 100% for species showing a distribution entirely in mountains). We defined a threshold (*T*) for species to be considered as montane, which refers to the minimum proportion of each species’ distribution overlapping with mountains. For example, a threshold of *T*=0.5 means that we considered a species to be montane when at least 50% of its distribution overlapped with mountains. To evaluate the sensitivity of our results to a certain threshold *T*, we performed all analyses using three different thresholds, *T*=0.3, *T*=0.5, and *T*=0.7. We considered that *T*=0.5 reflected the mean number of species in mountains and that using *T*=0.3 and *T*=0.7 reflected the range of standard error around this number (*T*=0.5±0.2, Fig. 1, Table 1, SI Appendix, Fig. S1 and Table S1). When using elevation-based definition of mountains, we further evaluated the sensitivity of this analysis to three different elevation limits (500, 700, and 1000 masl). As expected, we found that species richness in mountains decreased as the elevation threshold (*T*) increased; however, mountain area also diminished such that estimates of species per unit area did not show significant differences when changing elevation thresholds (SI Appendix, Fig. S8).

### Patterns of species richness along latitudinal gradients

To assess patterns of species richness along latitudinal gradients when controlling for area, we evaluated the relationship between species richness and latitude in either mountains and lowlands for birds, mammals and amphibians. We also performed the same analysis by summing the richness of the three groups simultaneously. Based on the rasterized maps of the distribution of the 21,741 vertebrate species, we generated a global map of species richness using a grid of 10 km x 10 km (around eight million pixels). To avoid spatial autocorrelation, we implemented a Monte Carlo procedure where we generated a random distribution of pixels where, for each of 1000 iterations, we randomly selected a sample of 1000 pixels located in mountains and 1000 pixels in lowlands. Because the area covered by land at northern latitudes is greater than in the tropics or at southern latitudes and corresponds mainly to lowland areas (SI Appendix, Fig. S9), we controlled for differences in area between tropical and temperate zones. To do so, we forced each subset of 1000 pixels to contain 500 pixels from tropical latitudes (below 23° latitude) and 500 pixels from temperate latitudes (above 23° latitude). We performed this procedure for mountain layers based on elevation and ruggedness. Finally, based on these subsamples, we ran a linear regression predicting local species richness against latitude and extracted the slope (β) and the coefficient of determination (R^2^) for mountains and lowlands separately. We compared the distributions of these two parameters between mountain and lowlands using an ANOVA (N = 1000). In addition, we tested for bias considering groups of species with different range size (from narrow-ranging to wide-ranging species), for which we divided the complete species pool in four 25% quantiles based on their distribution size, and performed the same analyses for each quantile (SI Appendix, Fig. S5). Finally, we also tested by differences among the main mountain ranges separated by continent that may exhibit region-specific patterns. For this, we ran the same regression analyses by dividing the global species pool in species in the same four quantiles but separating them in species occurring in the New World or the Old World (Fig. 2, SI Appendix, Fig. S6).

## Acknowledgments

We thank Carlos Guarnizo, Brian Smith, Daniel Cadena, and Jon Merwin for reading and making helpful comments to our manuscript. To BirdLife International and International Union for Conservation of Nature (IUCN) for providing the species distributions maps used in all analyses. Also, we thank BIOS (Centro De Bioinformática y Biología Computacional) for supporting us with the high computational requirements in its cluster server. Finally, we are grateful to Robert Ricklefs, Dolph Schluter, and the Laboratorio de Biología Evolutiva de Vertebrados from the Universidad de los Andes for valuable and productive discussions. The Instituto de Investigación de Recursos Biológicos Alexander von Humboldt supported this research.

## Supplementary information

**Fig. S1.**
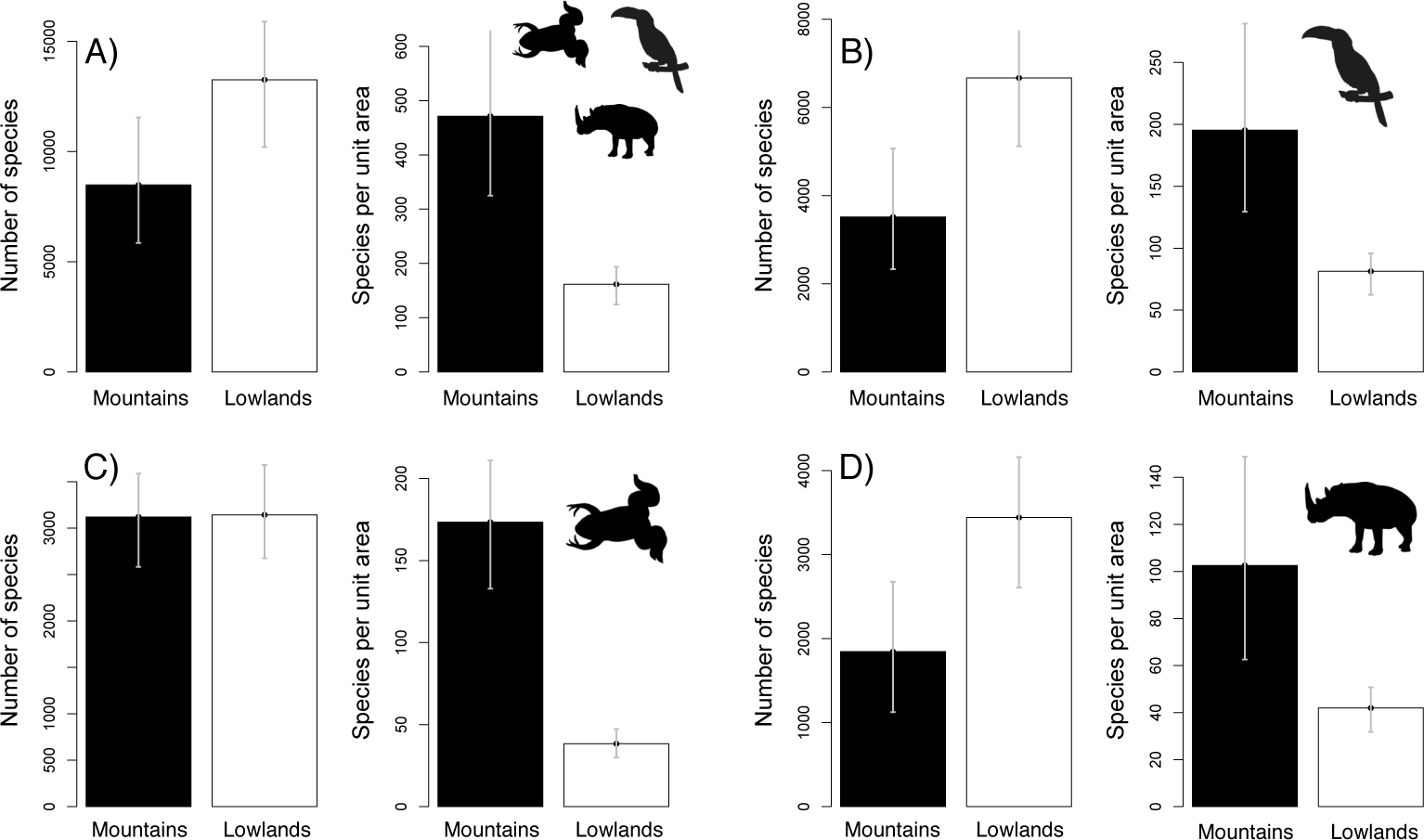
Differences in the contribution of mountains (black bars) and lowlands (white bars) to the total species richness of terrestrial vertebrates (global gamma-diversity) based on a ruggedness criterion to distinguish mountains from lowlands. The bars represent the total number of species (gamma diversity) and the number of species per unit area for all vertebrates (A), birds (B), amphibians (C), and mammals (D). In each panel, bar height corresponds to estimations using a *T*=0.5 and intervals correspond to the values generated based on *T*=70% and *T*= 30% (*T*= 50±20%).

**Fig. S2.**
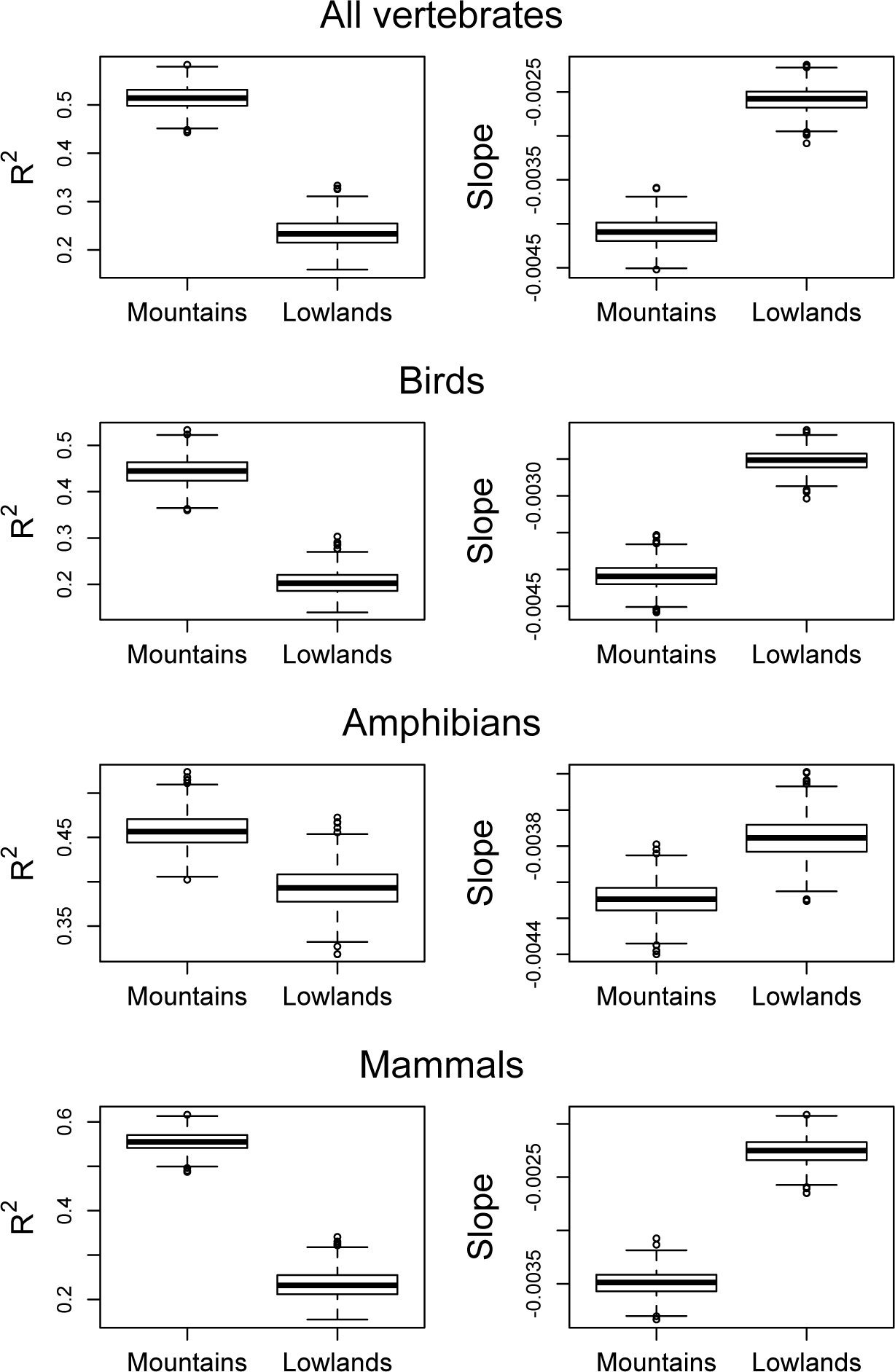
Boxplot illustrating differences in the coefficient of determination (*R*^2^) and the slope (β) of the linear regression predicting species richness against latitude based on a Monte Carlo procedure (1000 iterations) for mountains and lowlands, and using an elevation-based definition of mountain for all vertebrates (A), birds (B), amphibians (C) and mammals (D).

**Fig. S3.**
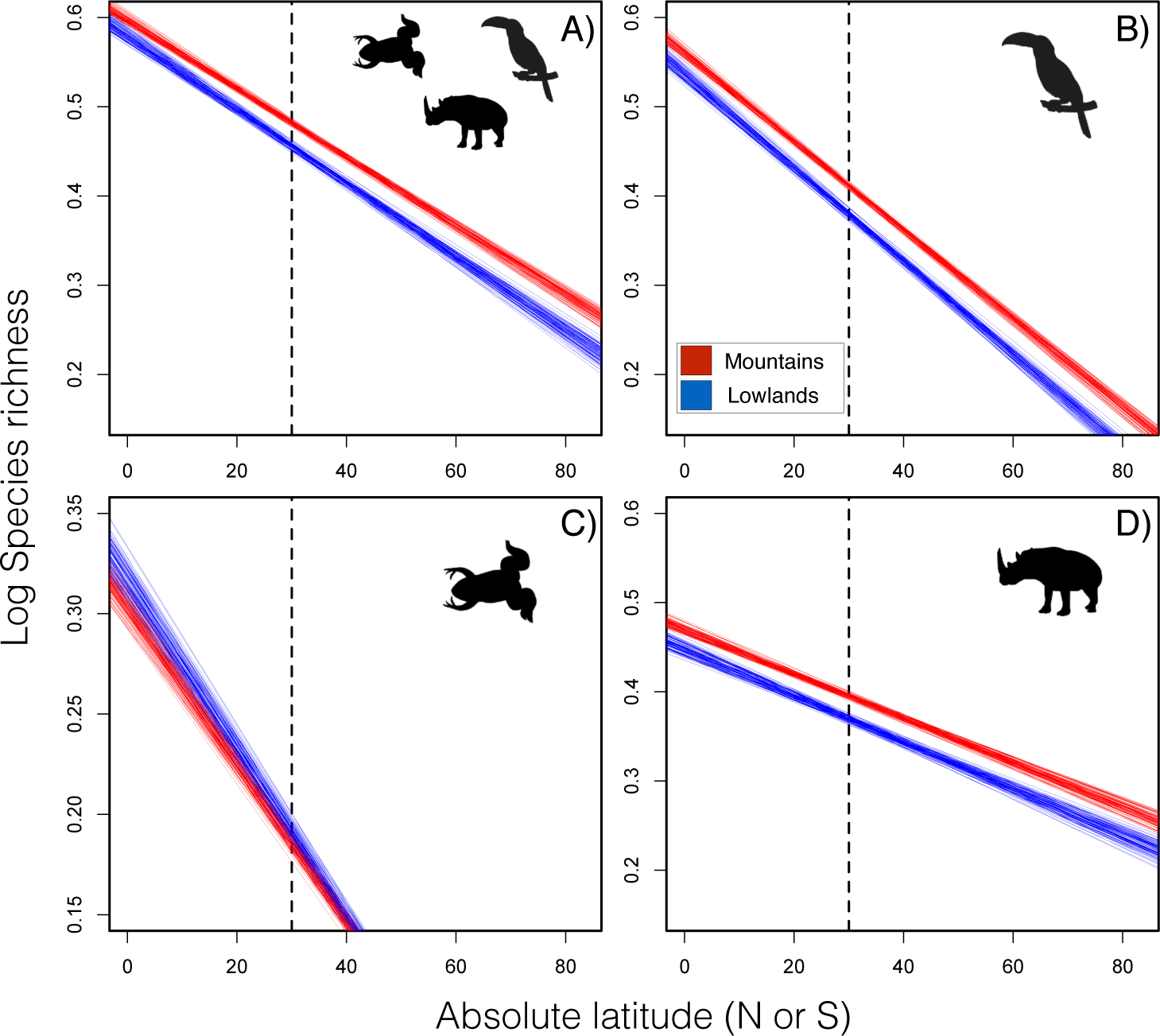
Patterns of species richness along the latitudinal gradient for mountains (red) and lowlands (blue) controlling by area, using a ruggedness criterion to limit mountains for all vertebrates (A), birds (B) amphibians (C), and mammals (D). Trend lines were extracted from 100 linear models based on a random sample of 1000 pixels for mountains and lowlands separately. The dotted black line represents 30° latitude, which represents the subtropical-temperate limit.

**Fig. S4.**
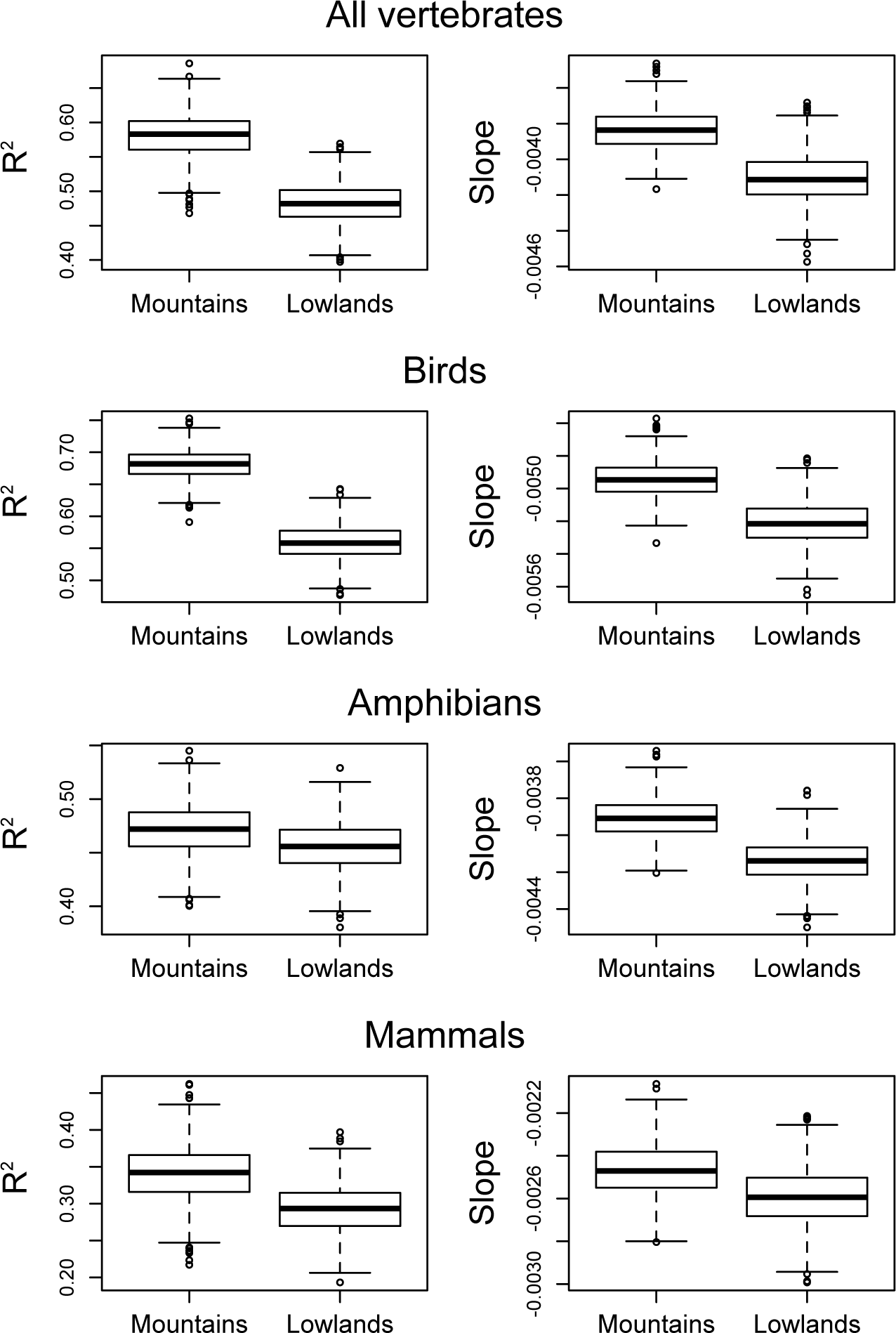
Boxplot illustrating differences in the coefficient of determination (*R*^2^) and the slope (β) of the linear regression predicting species richness against latitude based on a Monte Carlo procedure (1000 iterations) for mountains and lowlands, and using an ruggedness-based definition of mountain for all vertebrates (A), birds (B), amphibians (C) and mammals (D).

**Fig. S5.**
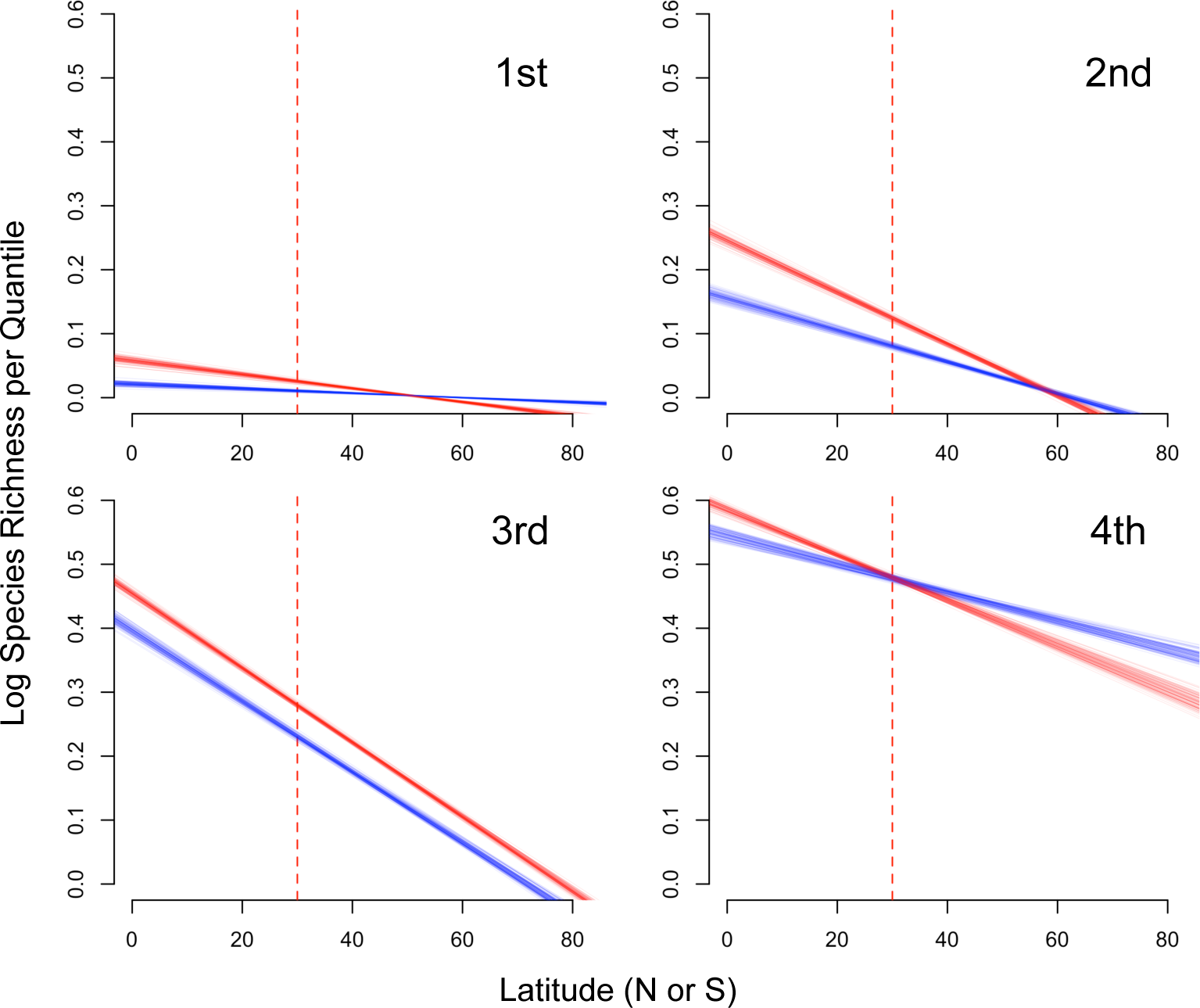
Species richness patterns by range size quartiles of terrestrial vertebrates richness (birds, amphibians, and mammals) along the latitudinal gradient for mountains (red) and lowlands (blue), controlling by area and using an elevation criterion to limit mountains. Each panel corresponds to a group of species divided in four quartiles according to their distribution range size, from the most restricted range species (1^st^, 0-25%) to the most widespread range species (4^th^, 75-100%). Trend lines were extracted from 100 linear models based on a random sample of 1000 pixels for mountains and lowlands separately. The dotted black line represents 30° latitude, which represents the subtropical-temperate limit.

**Fig. S6.**
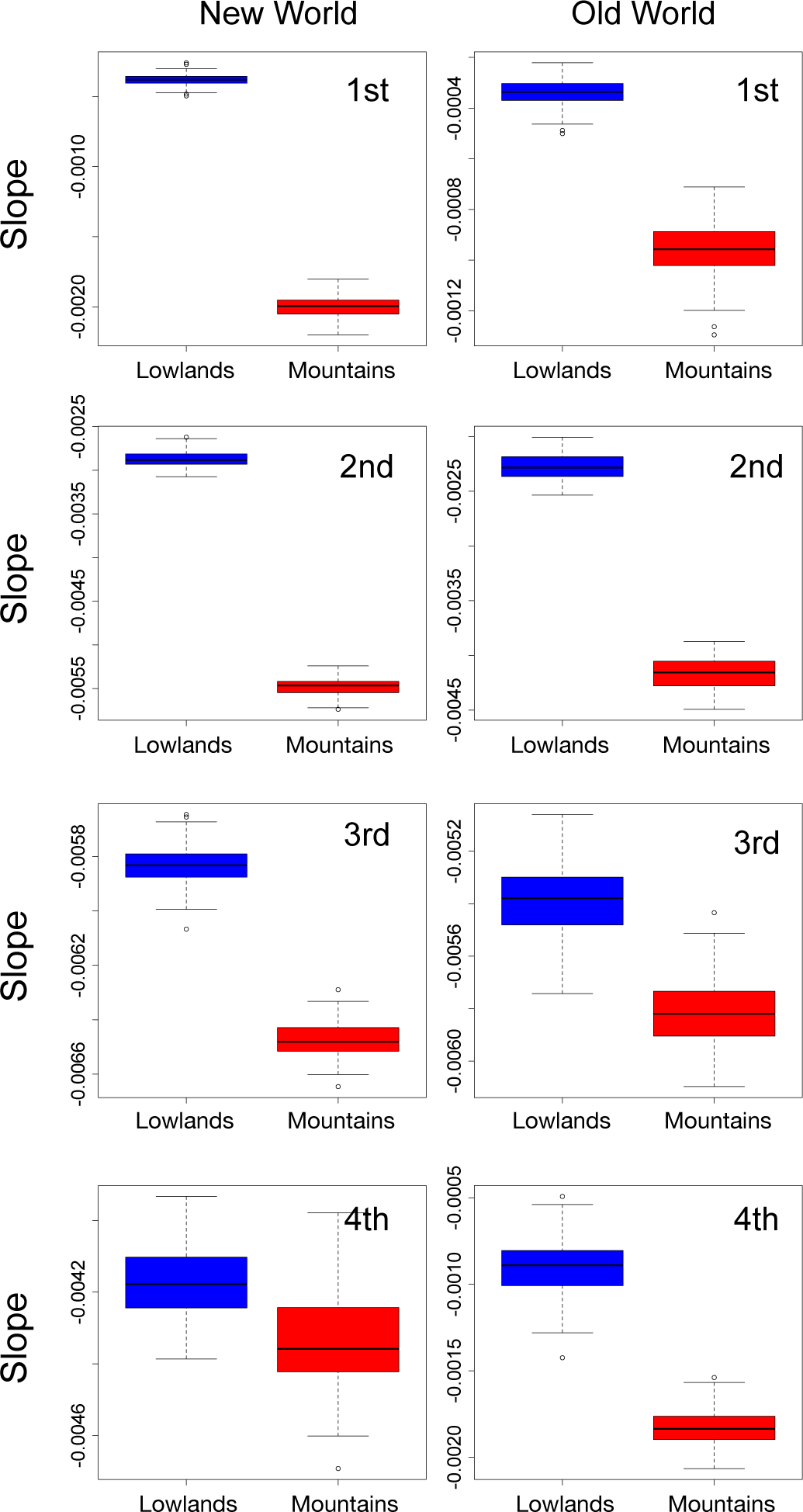
Boxplots of range size quantiles comparisons (four quantiles of 25%), illustrating differences in slopes (β) from linear regression predicting species richness against latitude between mountains (red) and lowlands (blue) based on a Monte Carlo procedure (1000 iterations). Analyses were performed separating pixels falling in the New World and Old World and using an elevation-based definition of mountain for all vertebrates. In all cases, significant differences were detected.

**Fig. S7.**
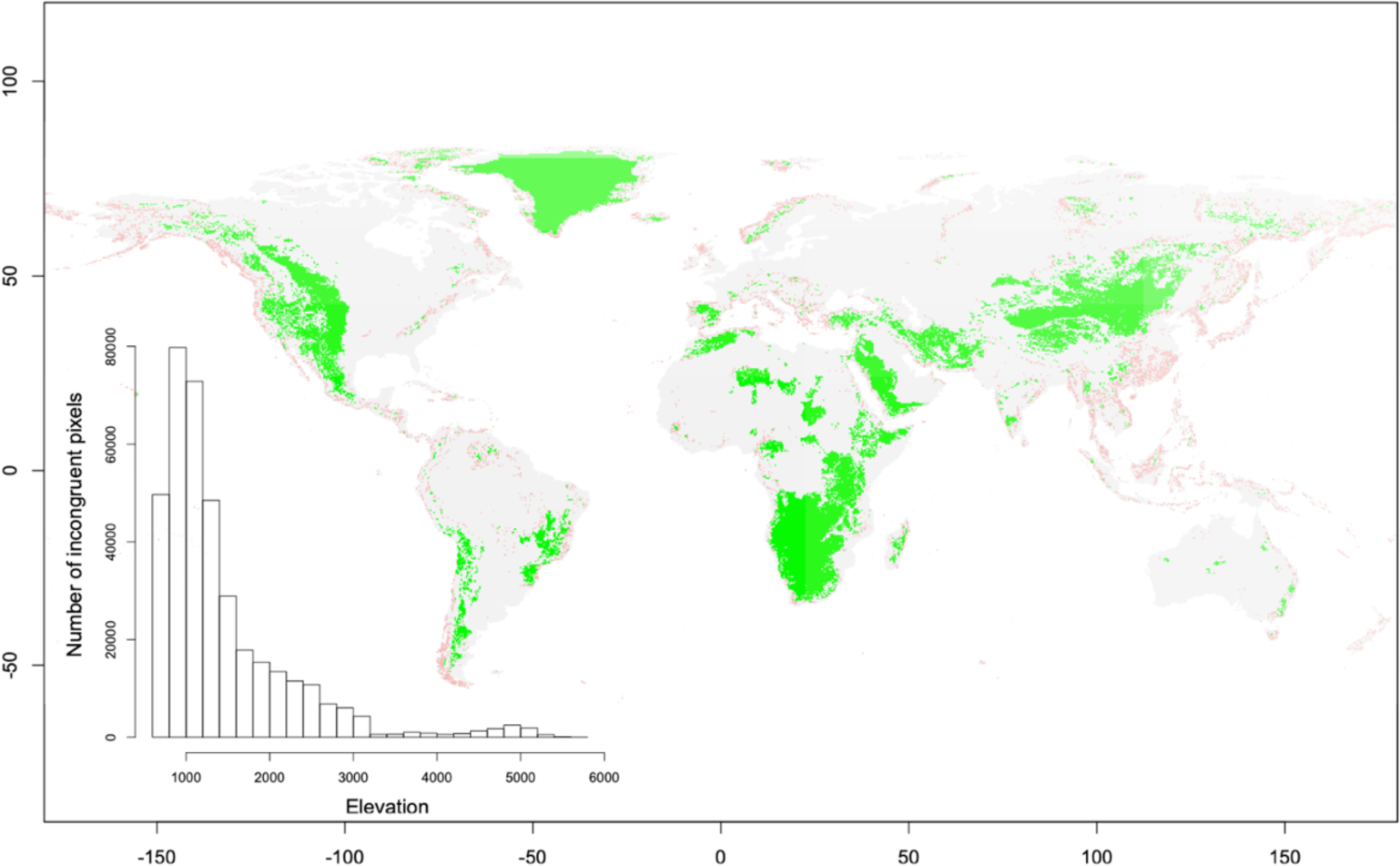
Geographical distribution of discrepant pixels between the two mountain definitions (Elevation-700 m and Ruggedness). Red pixels represent areas defined as mountains based on the Ruggedness raster layer, but not by the Elevation-700 raster layer. On the contrary, green pixels are these defined as mountains using the Elevation-700 raster layer, but not by the Ruggedness raster layer. The majority of discrepant pixels are located in middle elevations, between 500 and 2000 meters of elevation.

**Fig. S8.**
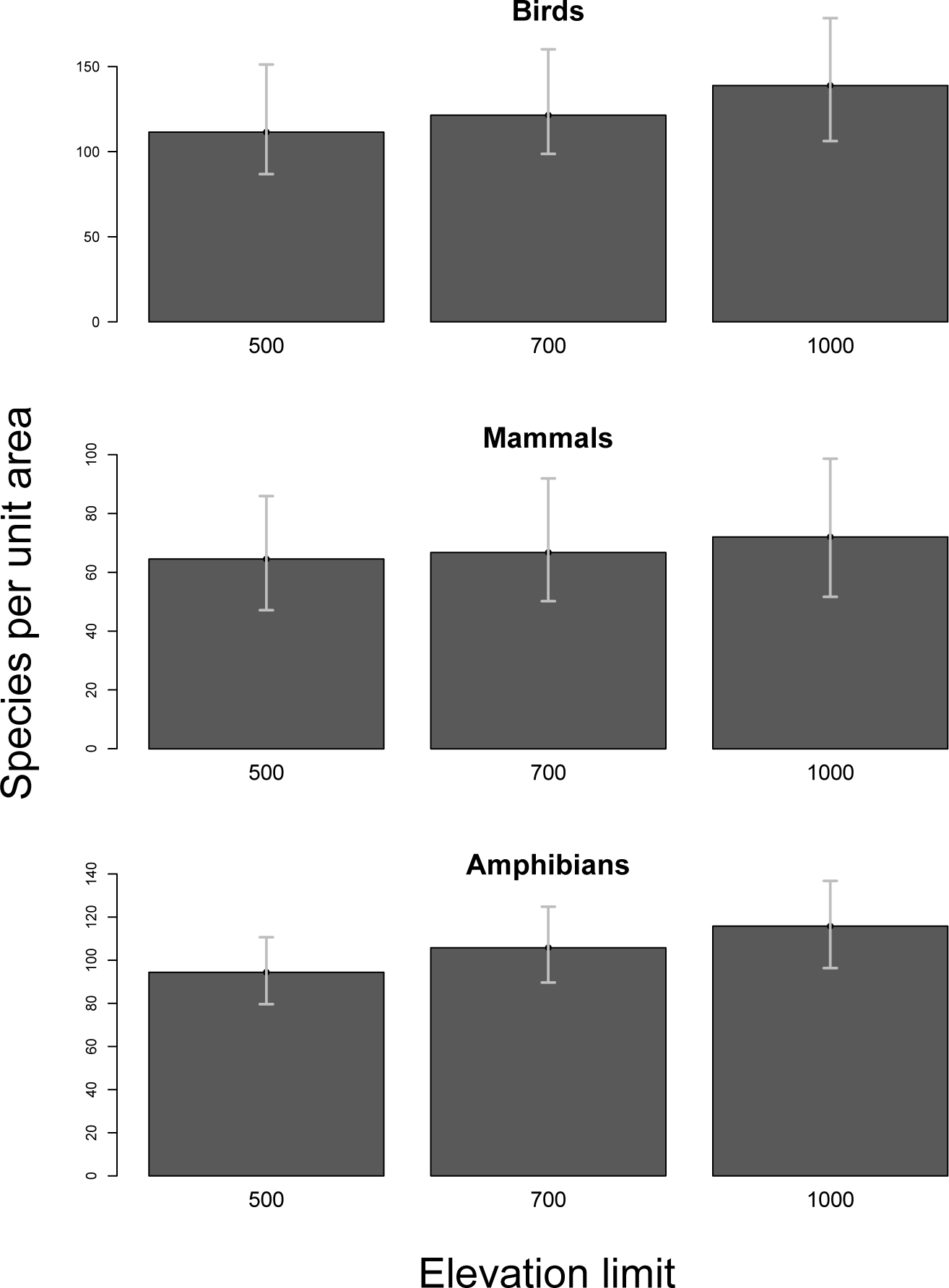
Bar plot of estimates of species per unit area for mountains using different elevation limits. Bar height corresponds to estimations using a *T*=0.5 and intervals correspond to values generated varying *T* (*T*= 0.5±0.2%, *T*=0.7% and *T*= 0.3%). Similar estimates are observed among the three elevation limits.

**Fig. S9.**
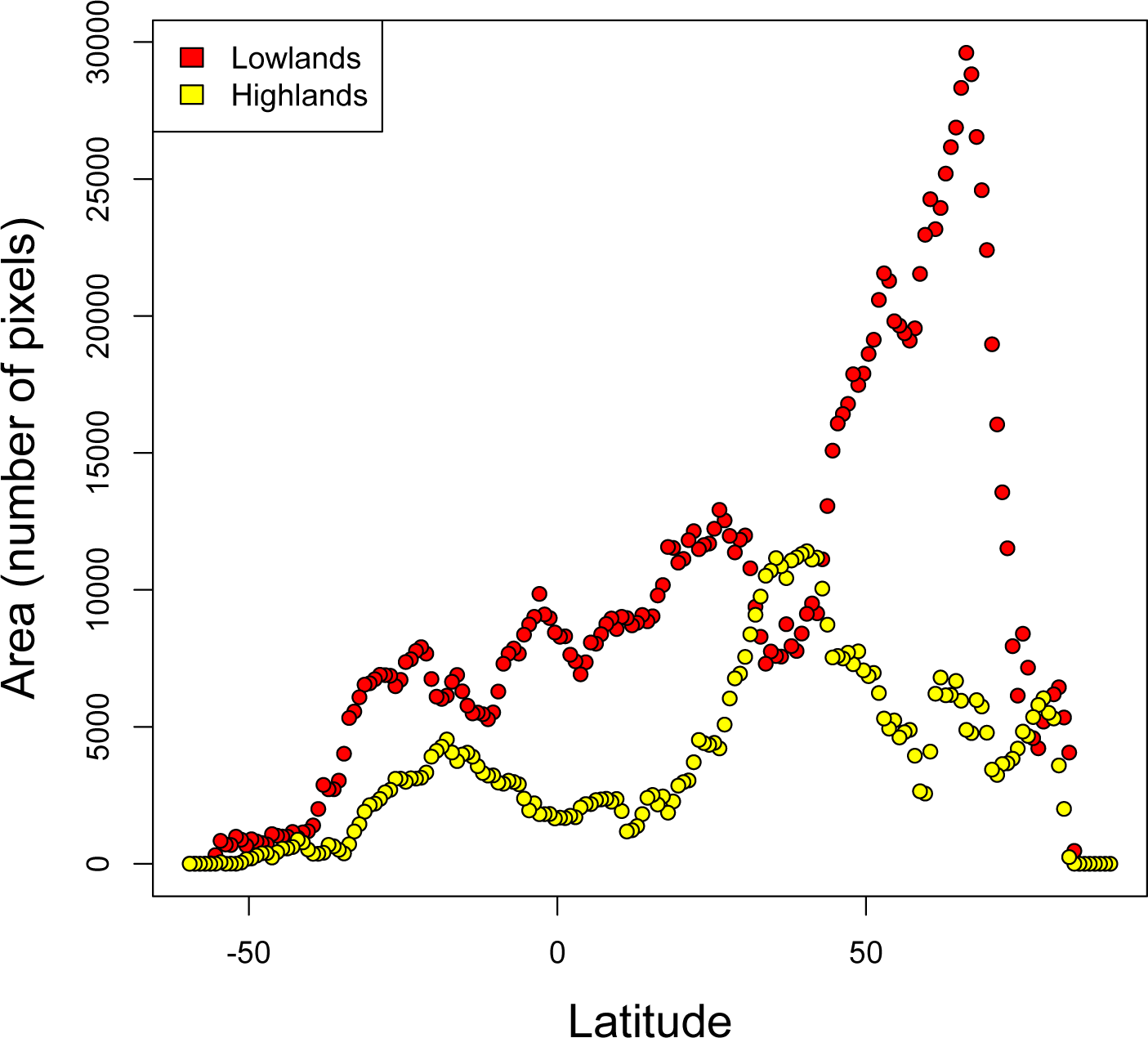
Estimation of area per latitudinal bands of 0.5° in mountains (yellow points) and lowlands (red points) as a function of latitude.

**Table S1.** Estimation of mountain and lowland contribution to the total species richness of terrestrial vertebrates (global diversity). Values are given for different thresholds (*T*) for defining a mountain species (0.5 ±0.2%), and based on a ruggedness-based mountain definition. Reported are the number of species in mountains and lowlands and the respective percentage is given in brackets.

| Group | Proportion of mountain distribution ($T$) | Mountain species | Lowland species | Mountain species per unit area | Lowland species per unit area |
| --- | --- | --- | --- | --- | --- |
| All groups | 0.3 | 11541 (53%) | 10195 (47%) | 641.17 | 124.33 |
|  | 0.5 | 8483 (39%) | 13253 (61%) | 471.28 | 161.62 |
|  | 0.7 | 5848 (27%) | 15888 (73%) | 324.89 | 193.76 |
| Birds | 0.3 | 5066 (50%) | 5118 (50%) | 281.44 | 62.41 |
|  | 0.5 | 3516 (35%) | 6668 (65%) | 195.33 | 81.32 |
|  | 0.7 | 2331 (23%) | 7853 (77%) | 129.50 | 95.77 |
| Amphibians | 0.3 | 3797 (61%) | 2467 (39%) | 210.94 | 30.09 |
|  | 0.5 | 3121 (50%) | 3143 (50%) | 173.39 | 38.33 |
|  | 0.7 | 2391 (38%) | 3873 (62%) | 132.83 | 47.23 |
| Mammals | 0.3 | 2678 (51%) | 2610 (49%) | 148.78 | 31.83 |
|  | 0.5 | 1846 (35%) | 3442 (65%) | 102.56 | 41.98 |
|  | 0.7 | 1126 (21%) | 4162 (79%) | 62.56 | 50.76 |

**Table S2.** Mean value and 95% confidence interval (in brackets) of the coefficient of determination (*R*^2^) and the slope (b) of the linear model predicting species richness against latitude for mountains and lowlands using the elevation-based definition of mountains.

| | Coefficient of determination ($R^2$) | | Slope ($\beta$) | |
| --- | --- | --- | --- | --- |
|  | Mountains | Lowlands | Mountains | Lowlands |
| All groups | 0.51 (0.47, 0.56) | 0.24 (0.18, 0.30) | -0.0041 (-0.0044, -0.0038) | -0.0026 (-0.0029, -0.0023) |
| Birds | 0.44, (0.38, 0.50) | 0.20 (0.15, 0.26) | -0.0041 (-0.0044, -0.0037) | -0.0025 (-0.0028, -0.0022) |
| Amphibians | 0.46 (0.42, 0.50) | 0.39 (0.35, 0.44) | -0.0041 (-0.0043, -0.0039) | -0.0038 (-0.0040, -0.0035) |
| Mammals | 0.56 (0.51, 0.60) | 0.23 (0.17, 0.29) | -0.0035 (-0.0037, -0.0033) | -0.0023 (-0.0025, -0.0020) |

**Table S3.** Mean value and 95% confidence interval (in brackets) of the coefficient of determination (*R*^2^) and the slope (b) of the linear model predicting species richness against latitude for mountains and lowlands using the ruggedness-based definition of mountains.

| | Coefficient of determination ($R^2$) | | Slope ($\beta$) | |
| --- | --- | --- | --- | --- |
|  | Mountains | Lowlands | Mountains | Lowlands |
| All groups | 0.58 (0.52, 0.64) | 0.48 (0.43, 0.54) | -0.0038 (-0.0040, -0.0036) | -0.0041 (-0.0043, -0.0039) |
| Birds | 0.68 (0.63, 0.72) | 0.56 (0.51, 0.61) | -0.0049 (-0.0052, -0.0047) | -0.0052 (-0.0054, -0.0049) |
| Amphibians | 0.47 (0.43, 0.52) | 0.46 (0.41, 0.50) | -0.0039 (-0.0041, -0.0037) | -0.0041 (-0.0044, -0.0039) |
| Mammals | 0.34 (0.27, 0.41) | 0.29 (0.23, 0.36) | -0.0025 (-0.0027, -0.0022) | -0.0026 (-0.0028, -0.0023) |

## Notes

### Competing Interest Statement

The authors have declared no competing interest.

